# Patterns of genetic variability in genomic regions with low rates of recombination

**DOI:** 10.1101/739888

**Authors:** Hannes Becher, Benjamin C. Jackson, Brian Charlesworth

**Affiliations:** Institute of Evolutionary Biology, School of Biological Sciences, University of Edinburgh, Charlotte Auerbach Road, Edinburgh EH9 3FL

## Abstract

Surveys of DNA sequence variation have shown that the level of genetic variability in a genomic region is often strongly positively correlated with its rate of crossing over (CO) [1–3]. This pattern is caused by selection acting on linked sites, which reduces genetic variability and can also cause the frequency distribution of segregating variants to contain more rare variants than expected without selection (skew). These effects of selection may involve the spread of beneficial mutations (selective sweeps, SSWs), the elimination of deleterious mutations (background selection, BGS) or both together, and are expected to be stronger with lower rates of crossing over [1–3]. However, in a recent study of human populations, the skew was reduced in the lowest CO regions compared with regions with somewhat higher CO rates [4]. A similar pattern is seen in the population genomic studies of *Drosophila simulans* described here. We propose an explanation for this paradoxical observation, and validate it using computer simulations. This explanation is based on the finding that partially recessive, linked deleterious mutations can increase rather than reduce neutral variability when the product of the effective population size (*N*_*e*_) and the selection coefficient against homozygous carriers of mutations (*s*) is ≤ 1, i.e. there is associative overdominance (AOD) rather than BGS [5]. We show that AOD can operate in a genomic region with a low rate of CO, opening up a new perspective on how selection affects patterns of variability at linked sites.

## Results and Discussion

### Diversity Statistics in Relation to CO Rates in *Drosophila simulans*

The top two panels of Figure 1 show the relations between the rate of crossing over and mean pairwise diversity at four-fold degenerate nucleotide sites (*π*_4_), for two populations of *Drosophila simulans*. Consistent with previous studies of many species [1–3, 6], there is a significant positive relation between CO rate and nucleotide site diversity for both X chromosome (X) and autosomes (A). As reported previously [7], X has much lower diversity than A for all bins of CO rates. Diversity is higher for the Madagascan than the Kenyan population, consistent with the latter having been founded as a relatively small population by flies descended from the putatively ancestral Madagascan population [7].

**Figure 1.**
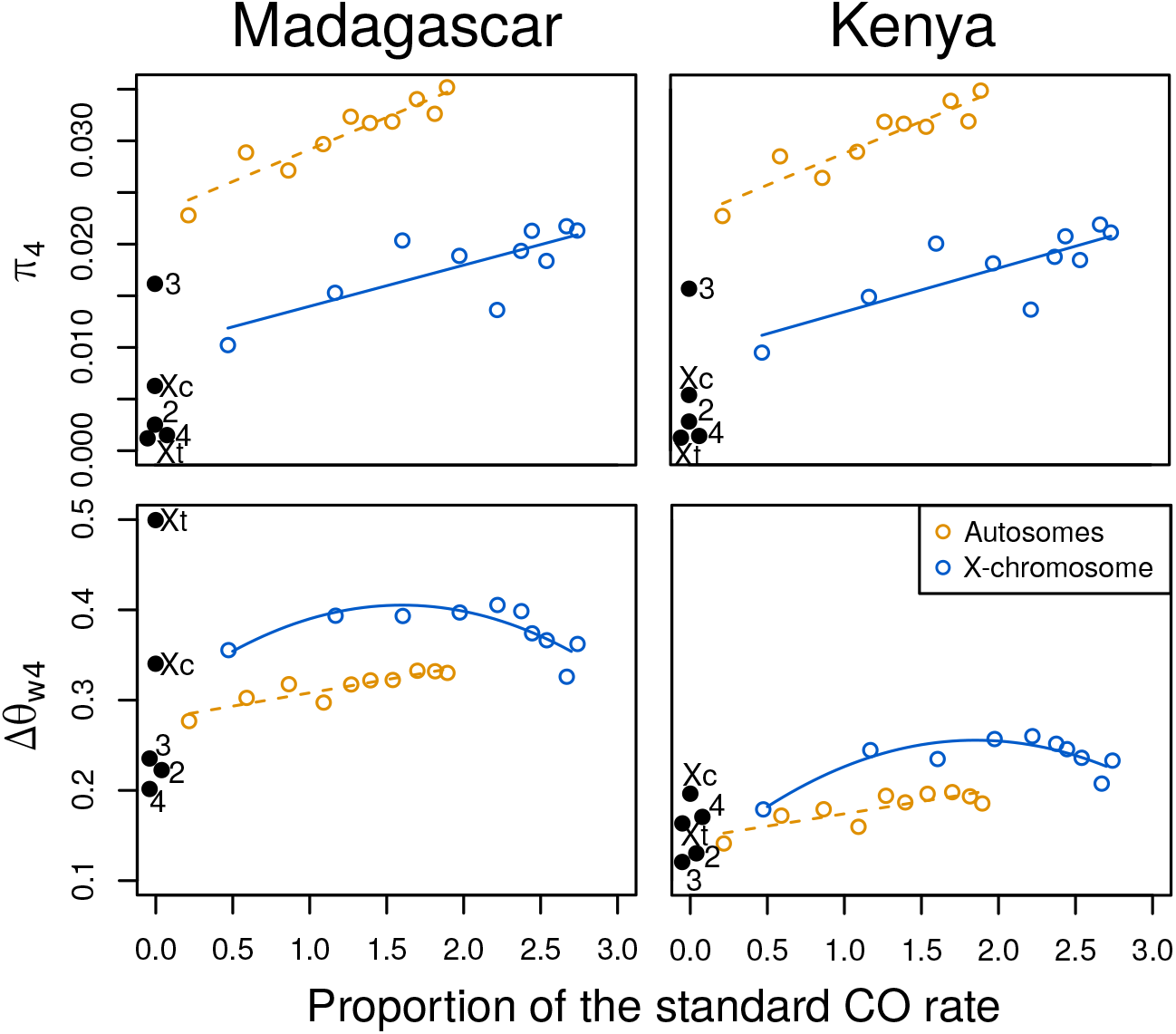
Genetic Diversity Statistics at Four-fold Degenerate Sites in Two Populations of *Drosophila simulans* in Relation to the Rate of Crossing Over. All panels share the same X axis, the mean proportion of the standard crossover (CO) rate per megabase in homologous regions of *Drosophila melanogaster* for each group of genes. Estimates of the pairwise nucleotide diversity (π_4_) for each group are shown in the top panels, and estimates of a measure of the skew of the site-frequency spectrum (Δθ_w4_) in the bottom panels. These statistics were computed for each gene and then binned by CO rate, after weighting by the number of four-fold sites per gene in each bin. Autosomal bins are shown in orange and X chromosomal bins in blue. The solid lines indicate significant regression fits for the X chromosomal values, with the following significance levels for the quadratic terms – Madagascar π_4_: *p* = 0.00960; Madagascar Δθ_w4_: *p* = 0.0083; Kenya π_4_: *p* = 0.00598; Kenya Δθ_w4_: *p* = 0.00262. The dashed lines indicate linear regression fits for the autosomal bins. The significance levels from Spearman rank correlations were as follows – Madagascar π_4_: ρ = 0.939, *p* < 2.2 × 10^−16^; Madagascar Δθ_w4_: ρ = 0.891, *p* = 0.00138; Kenya π_4_: ρ = 0.927, *p* = 0.000130; Kenya Δθ_*w*4_: ρ = 0.685, *p* = 0.0351). Non-CO regions are shown as black dots. These are spread out for better visibility and are labelled with their identities; Xc is the centromeric region of the X and Xt is its telomeric region; 2 and 3 are centromeric regions of chromosomes 2 and 3, respectively; 4 is the dot chromosome.

The bottom panels of Figure 1 show the relations between CO rate and a measure of the skew of the site frequency spectrum (SFS) towards rare variants, Δθ_w4_ = 1 − π_4_ / θ_w4_, where π_4_ and θ_w4_ are the estimates of diversity based on the mean pairwise difference between alleles [8] and the number of segregating sites [9], respectively. Other measures of skew behave similarly (Supplemental Figure S1). As was also found in a Rwandan population of *D. melanogaster* [10], there is much greater skew on X than A, although the absolute level of skew for both X and A is much higher than in *D. melanogaster*, suggesting a more intense recent population expansion in *D. simulans*. The skew is also larger in the Madagascan than the Kenyan population, consistent with the latter having experienced a bottleneck in population size or a slower rate of population expansion. The parameters of the population expansion for the Madagascan population were estimated by [11].

The most surprising feature of these results is that there is a nearly monotonic increase of Δθ_w4_ with CO rate in the autosomal CO regions. The overall autosomal values of Δθ_w4_ for non-CO regions are 0.22 and 0.13 for Madagascar and Kenya, respectively. A meta-analysis of 10 independent autosomal non-CO regions from six *Drosophila* species (Supplemental Table S1) gave a mean of 0.18, with s.e. 0.04. The pattern is more complex for the X in *D. simulans* – there is a significant quadratic relation between Δθ_w4_ and CO rate, with an initial increase followed by a decrease at the highest CO rates. The non-CO regions of the X show values of Δθ_w4_ comparable to the CO regions, with the exception of the telomere in the Madagascan population, which has a value of 0.44, possibly reflecting a recent selective sweep, consistent with its very low π_4_ value.

If the skew in the SFS were in part caused by selective sweeps (SSWs), it would be expected to decrease, not increase, with CO rate, as was found in computer simulations with parameter values that are realistic for *Drosophila* [12]. If background selection (BGS) also contributes, these simulation results showed that it should only produce a strong effect on skew in the non-CO regions, and cause these to have higher skews than regions with CO. Indeed, the simulations predicted higher skews in the non-CO regions than the observed autosomal means presented above: Δθ_w4_ was 0.24 for autosomal simulations with BGS and SSWs, and 0.21 with BGS alone. Given that demographic effects probably contribute substantially to the skew in the SFS in *D. simulans*, especially in Madagascar [11], an even higher proportion of singletons in the non-CO regions is expected.

We will address elsewhere the question of relations between the level of skew of the SFS towards low frequency variants and the CO rate outside the non-CO regions, and simply note that it probably reflects the joint effects of SSWs and population size changes, which vary among genomic regions with different effective population sizes (*N*_*e*_) associated with different CO rates, so that low *N*_*e*_ can reduce the effect on skew of a recent population expansion [10]. We focus here on the fact that the level of skew in the non-CO regions is smaller than expected with BGS and/or SSWs.

### Simulation Results and an Explanation for Low Skew

The simulations in [12] assumed a dominance coefficient (*h*) of 0.5 for deleterious mutations, where the fitness of the heterozygous carriers of a deleterious mutation is reduced by *hs* relative to wild-type, with homozygotes suffering a reduction of *s*. However, studies of the effects of deleterious mutations on fitness components [13–15] suggest that there is likely to be partial recessivity of most slightly deleterious mutations, with 0 < *h* < 0.5. Because selection against rare deleterious mutations in randomly mating populations acts mainly on their heterozygous carriers [16], the value of *h* does not greatly affect the fate of rare deleterious mutations with similar values of *hs*, so that *h* = 0.5 was used previously for convenience.

Empirical estimates of the parameters of the distribution of deleterious mutational effects on fitness (DFE) in *Drosophila* suggest that it has a large standard deviation relative to its mean [17–20]. Under a gamma distribution, the shape parameter for both nonsynonymous (NS) and UTR mutations has been estimated to be approximately 0.3 [20]. Such a wide distribution implies that there is a substantial proportion of selection coefficients against mutant homozygotes (*s*) that have *N*_*e*_*s* of the order of 1. For example, with *h* = 0.2, a shape parameter of 0.3 and 2*N*_*e*_*s* = 5000 (the value for NS sites in our simulations), the proportion of mutations with 2*N*_*e*_*s* ≤ 2.5 is 0.08.

This raises the possibility that a subset of partially recessive deleterious mutations could cause associative overdominance (AOD), whereby loss of variability at neutral sites is retarded by linkage disequilibrium (LD) between selected and linked neutral sites [21]. Such LD causes a correlation in homozygosity between sites, so that homozygotes for variants at the neutral sites are associated with reduced fitness compared to heterozygotes when deleterious mutations have some degree of recessivity [5]. Recent theoretical work has shown that partially recessive deleterious mutations can result in AOD when 2*N*_*e*_*s* is of the order of 2.5 [5], given sufficiently close linkage between neutral and selected sites, which is prevalent in low recombination genomic regions. By definition, AOD acts similarly to true overdominance, a form of balancing selection. Long-term balancing selection is known to increase the lengths of the internal branches of coalescent trees relative to the total tree size [22–25]; AOD should have a similar effect, causing the SFS to become biased towards more frequent variants than with strict neutrality. BGS without AOD has the opposite effect, especially in low CO regions where selective interference among deleterious mutations increases the skew towards rare variants [12, 26–30].

If the more abundant strongly selected mutations cause standard BGS effects in a low recombination region, but some weakly selected mutations cause AOD, the net outcome for a low CO region should be lower diversity and a site frequency spectrum (SFS) more skewed towards low frequency variants than with no selection, but with higher diversity and less skew than with standard BGS. This process could account for the relatively low skew towards low frequency variants in the non-CO regions of humans [4] and six *Drosophila* species (Supplemental Table S1).

We have investigated this possibility by simulations using a range of different dominance coefficients (see STAR Methods for details). Figure 2A shows results with *h* = 0.2 and *h* = 0.5, for a range of CO rates that are all ≤ the mean rate of 2 × 10^−8^ for female meiosis in *D. melanogaster* [31, 32]. With deleterious mutations alone, advantageous mutations alone, and with both deleterious and advantageous mutations, the mean value of neutral diversity increases as the CO rate increases, whereas the degree of skew towards low frequency variants decreases. In the presence of deleterious mutations, the degree of skew in very low CO rate regions is substantially lower with *h* = 0.2 than *h* = 0.5, both in the presence (0.16, 0.24) and absence of advantageous mutations (0.10, 0.21), and is not far from the observed mean autosomal value across *Drosophila* species of 0.18. In addition, neutral diversity with deleterious mutations is higher for *h* = 0.2 (0.0041) than *h* = 0.5 (0.0024) in non-CO regions. Figure 2B shows the behaviour of a non-CO region with a range of dominance coefficients, showing that low *h* greatly reduces the skew caused by BGS, and increases diversity. As was found previously [12], BGS is the main cause of reduced variability in the non-CO cases, reflecting the fact that the rate of sweeps is greatly reduced by BGS when CO is absent. These findings are consistent with the hypothesis that AOD (caused by the relatively small fraction of mutations that are weakly selected) reduces the effect of BGS when mutations are partially recessive.

**Figure 2.**
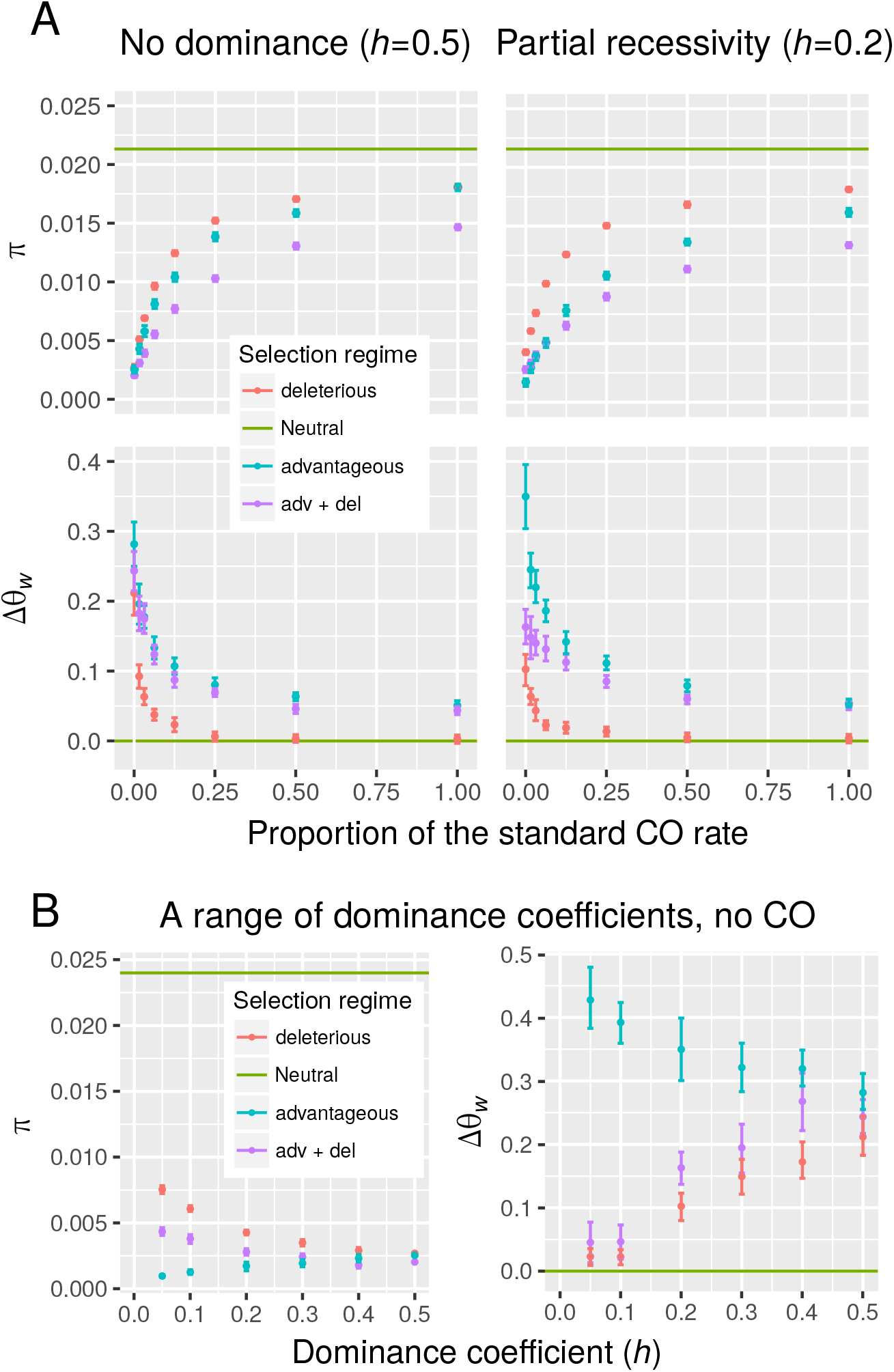
Expectations for the Patterns of Genetic Diversity Statistics from Forwards-in-Time Simulations. (A) The pairwise diversities at simulated autosomal synonymous sites (π, top row) and a measure of the distortion of the site-frequency spectrum (Δθ_w_, bottom row) for different rates of crossing over, expressed as a proportion of the mean sex-averaged rate for autosomes in *D. melanogaster*. Gene conversion was allowed with the standard parameter values for *D. melanogaster*, regardless of the CO rate. The theoretical expectations with no selection are given by the horizontal green lines. Simulations with selection against deleterious mutations alone are indicated as ‘deleterious’, simulations with advantageous mutations alone as ‘advantageous’, and those with both kinds of selection as ‘adv + del’. The results in the left-hand panels are for simulation runs with semi-dominant mutations (*h* = 0.5), and those in the right-hand panels are with partial recessivity (*h* = 0.2). Each data point shows the mean and 95% confidence intervals for 20 replicate simulations, for a group of 70 genes subject to selection (see STAR Methods for details of the parameters used in the simulations). (B) Simulated values of π and Δθ_w_ for a range of different dominance coefficients (*h*) without crossing over but with gene conversion occurring at the standard rate for *D. melanogaster*

### Linkage Disequilibrium Patterns and a Further Test for AOD

This hypothesis can be further tested by noting that that AOD increases the relative lengths of internal branches of the coalescent tree for a neutral site (see above). By analogy with the effects on LD of a recent population size change, this change in tree topology increases the magnitude of LD [33, 34]. Conversely, BGS with low CO rates has the opposite effect, since it increases the relative lengths of the external branches of coalescent trees, as noted above. We can use the simulated values of neutral diversity to estimate the corresponding value of *N*_*e*_ from the formula π = 4*N*_*e*_*u*, where *u* is the mutation rate per site) [35]. This can be used to determine the approximate expected value of *r*^2^, the squared correlation coefficient between the allelic states of pairs of neutral sites from equation (3) in [36]. As can be seen in Figure 3 (top left), the simulation values of median *r*^2^ for all pairs of neutral sites with deleterious mutations and without CO (but with gene conversion) are more than an order of magnitude lower than the neutral expectations, which are 0.200 with *h* = 0.2 and 0.209 with *h* = 0.5, indicating a strong influence of BGS. Nonetheless, the magnitude of LD with *h* = 0.2 is much greater than with *h* = 0.5 despite a higher level of diversity and hence *N*_*e*_ (note that low *N*_*e*_ is expected to increase *r*^2^).

**Figure 3.**
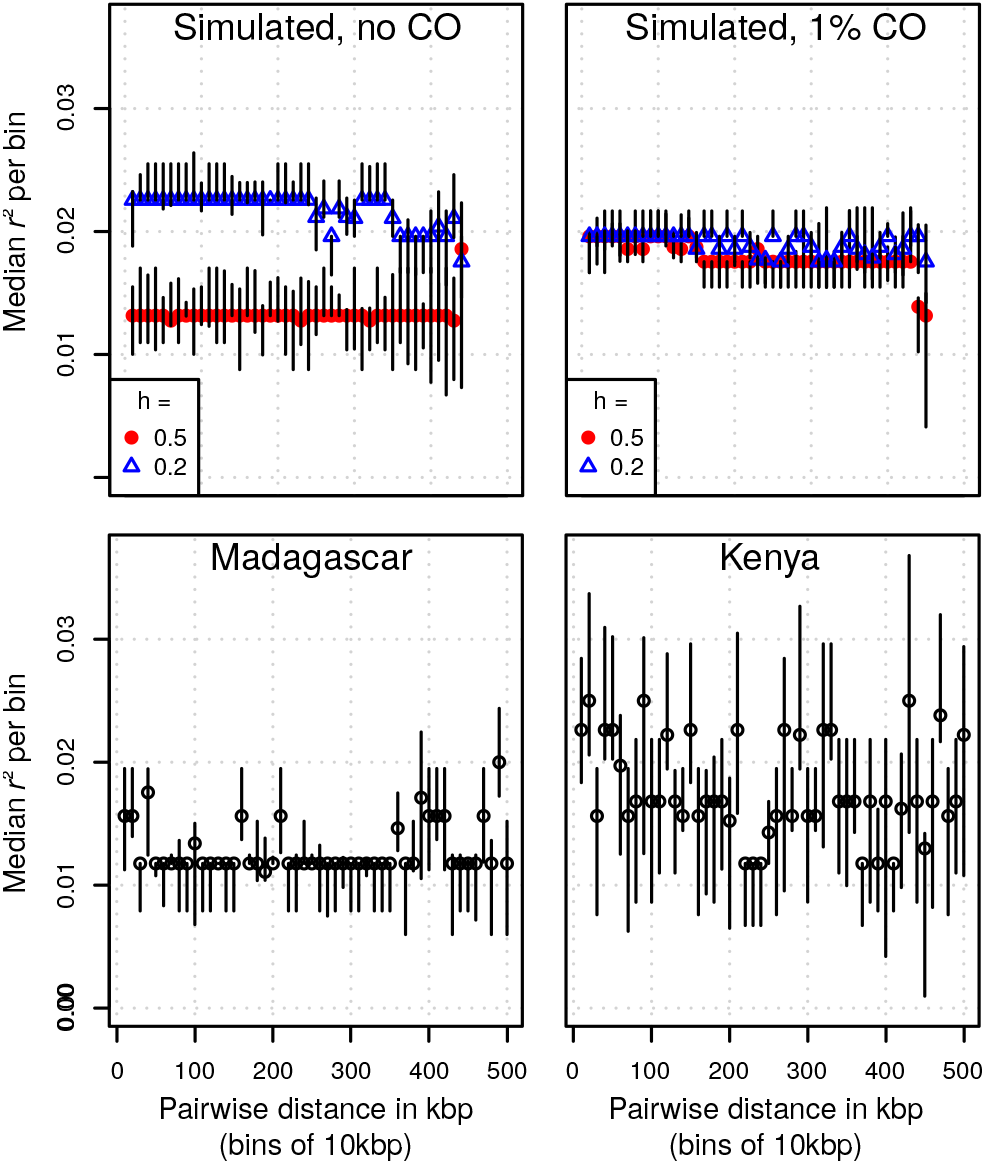
Linkage Disequilibrium (LD) as Measured by the Median of *r*^2^ Between Pairs of Sites. The top panels show simulation results for a diploid population size of *N* = 2500 with BGS and sweeps, with either no CO or CO at 1% of the standard rate. Red circles indicate runs without dominance (*h* = 0.5) and blue triangles indicate runs with partial recessivity (*h* = 0.2). Mantel tests carried out on individual simulation runs are only significant in the presence of CO. The bottom panels show the average values of the LD measure across bins of pairwise distances, between four-fold degenerate variant sites on chromosome four for the Madagascan and Kenyan populations. Mantel tests carried out on the (un-binned) data show that there is a significant decay of LD with increasing distance (Madagascar four-fold: 222 sites, Mantel statistic *R* = 0.061, *p* = 0.001; Madagascar zero-fold: 320 sites, *R* = 0.040, *p* = 0.001; Kenya four-fold: 187 sites, *R* = 0.026, *p* = 0.015; Kenya zero-fold: 267 sites, *R*= 0.027, *p* = 0.002).

The bottom panels of Figure 3 show plots of median *r*^2^ for 4-fold sites against physical location for 50 bins of distances between pairs of variants across genes on the non-CO chromosome four in the Madagascan and Kenyan populations. As is expected from bottleneck effects or a lower rate of population expansion, LD is markedly higher in Kenya than Madagascar, but the overall levels of LD are of similar magnitude to the simulation results, and are much lower than predicted from the estimates of *N*_*e*_ using π_4_ and the *D. melanogaster* mutation rate [37]. There is an indication that there is a low level of crossing over on this chromosome, from the significant negative relation between *r*^2^ and distance between pairs of sites, which is not observed in simulations with gene conversion alone.

It is, of course, possible that true balancing selection, rather than AOD, could contribute to these patterns. Supplemental Figure S2 shows plots of 4-fold and 0-fold diversities per gene along chromosome four, for the Madagascan and Kenya populations. There is one gene, FBgn0268752, which stands out as having unusually high diversities at both types of site in both populations. In the case of Madagascar, there are two intermediate-frequency nonsynonymous variants (at positions 40149 and 40648) (Supplemental Table S2), whereas Kenya is segregating for nonsynonymous variants only at 40648, possibly because of loss of variability after a bottleneck. The polymorphisms at sites 40149 and 40648 in Madagascar are not in LD with each other, whereas there are strong associations between one of the variants at 40149 and synonymous variants to its left (Supplemental Table S2). It is, therefore, possible that there is balancing selection acting at this locus, which appears to be unique to *D. simulans.* However, there is no evidence that the LD created by the 40149 variant extends outside this locus, so that it does not contribute to the general patterns described above.

Previous theoretical work on AOD due to deleterious, partially recessive mutations suggested that it is likely only to occur in small populations, where there is indeed evidence that the rate of loss of variability for molecular markers is slower than expected from the neutral value of *N*_*e*_ [5]. This is because AOD requires *N*_*e*_*s* to be of the order of 1, which is likely to be met by only a small minority of deleterious mutations in a large, randomly mating population. However, when *N*_*e*_ for neutral sites is greatly reduced by selection at linked sites, the extent of randomly generated LD between neutral and selected sites will be considerably enhanced, and LD is the driving force for AOD [5].

The simulation results shown in Figures 2 and 3 suggest that, with partially recessive mutations and the wide distribution of *s* values suggested by population genomic analyses of the effects of deleterious mutations, there is a noticeable effect of AOD on the SFS and extent of LD at neutral sites when CO rates are very low. This interpretation can be tested by comparing the results of simulations with a narrow versus wide range of selection coefficients for deleterious mutations, keeping the mean constant (Figure 4). In this case, diversity, skew and the magnitude of LD are independent of *h*; with the wide range of selection coefficients, diversity and LD decrease with *h*, but skew increases. This is what is expected if AOD is influencing the behaviour of neutral variants in low CO regions.

**Figure 4.**
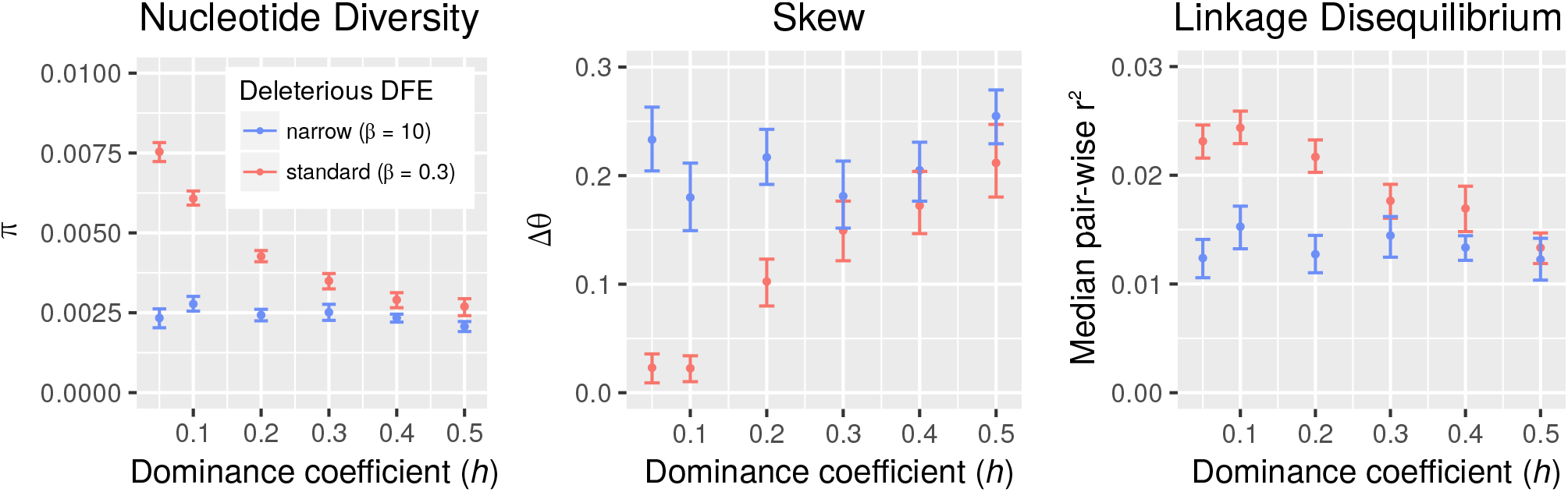
Weakly Deleterious Mutations are Required for AOD Effects on Variability. The panels show the results of simulations with deleterious mutations alone, in the absence of CO and a range of dominance coefficients, with the mean strength of selection being kept constant. Red and blue indicate two different distributions of deleterious fitness effects with the same mean but different shape parameters (β). The results with β = 0.3 (a wide distribution of selection coefficients), are the same as those shown previously, and are contrasted with the results for a narrow distribution (β = 10), where AOD is unlikely to occur.

The properties of genomes or genomic regions with low rates of genetic recombination are of considerable interest for a range of biological questions, including evolution and variation in bacteria [38, 39], asexual higher organisms [40, 41], organisms with high rates of self-fertilisation [42, 43], and Y and W chromosomes [44–47]. There is a general expectation that genetic diversity and molecular signatures of adaptive evolution should be greatly reduced in such systems, as a result of selection at linked sites. This reduction in adaptation is important for understanding phenomena such as the degeneration of Y chromosomes and the lack of evolutionary success of asexual and highly selfing species.

However, comparisons between different species with different modes of reproduction that influence the recombination rate are often made difficult by confounding factors, such as differences in the extent of colonisation events involving population size bottlenecks [42, 43]. The use of comparisons among genomic regions with different recombination rates largely removes this difficulty, and has revealed many of the expected results [2, 3, 6]. Here, we have shown that an unexpected pattern emerging from such comparisons suggests that the largely neglected process of associative overdominance may influence patterns of variability when recombination rates are very low. Because it requires partial recessivity of the fitness effects of deleterious mutations, and the opportunity for them to be present in both heterozygous and homozygous states, it will only operate in diploid or polyploid organisms with some degree of outbreeding. It will, therefore, not affect haploid, asexual or highly selfing organisms, or effectively haploid chromosomes such as Y and W. This suggests that we might expect to see differences in their patterns of variability compared with the non-CO regions of outbreeding diploid species.

## Supporting information

GZ-compressed archive of supplemental tables (XLSX)

## Author contributions

BC designed the project and contributed to the analyses of the data and simulations; HB conducted the simulations and contributed to the analyses of the population data and simulations; BCJ conducted the bioinformatic analyses of the raw sequencing data. All three authors contributed to the writing of the manuscript.

## Acknowledgements

We wish to thank Ching-Ho Chang and Amanda Larracuente for providing the sequence diversity data for the dot chromosome in *D. pseudoobscura* and *D. miranda*. We made use of the resources provided by the Edinburgh Compute and Data Facility (ECDF) (http://www.ecdf.ed.ac.uk/). This research was funded by a Leverhulme Trust grant (IRPG-2015-033) to BC.

## Supplemental Information for Becher, Jackson & Charlesworth

**Supplemental Figure S1.**
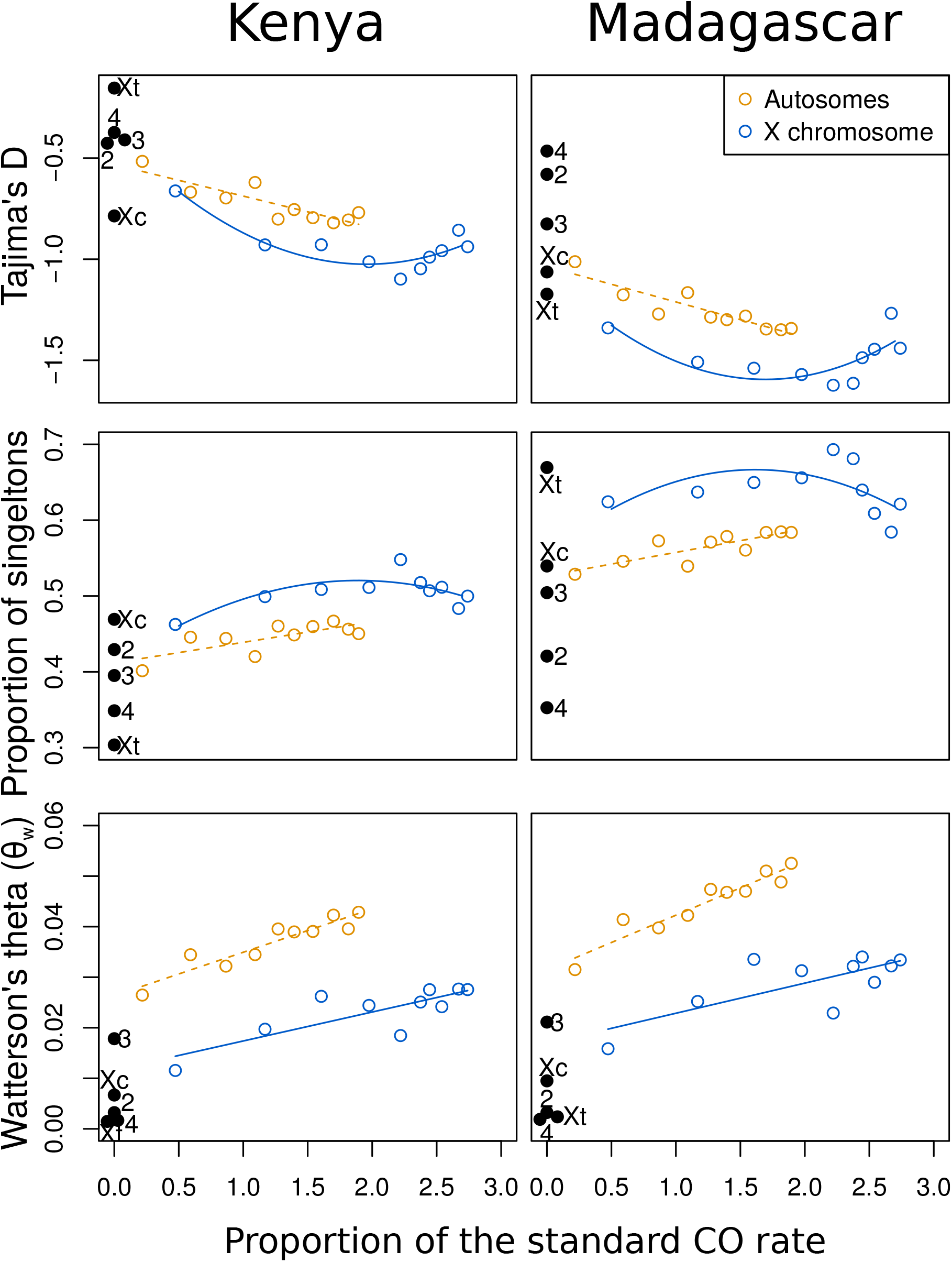
Related to figure 1. Additional statistics for the skew of the site-frequency spectrum and Watterson’s theta, an estimate of sequence diversity.

**Supplemental Figure S2.**
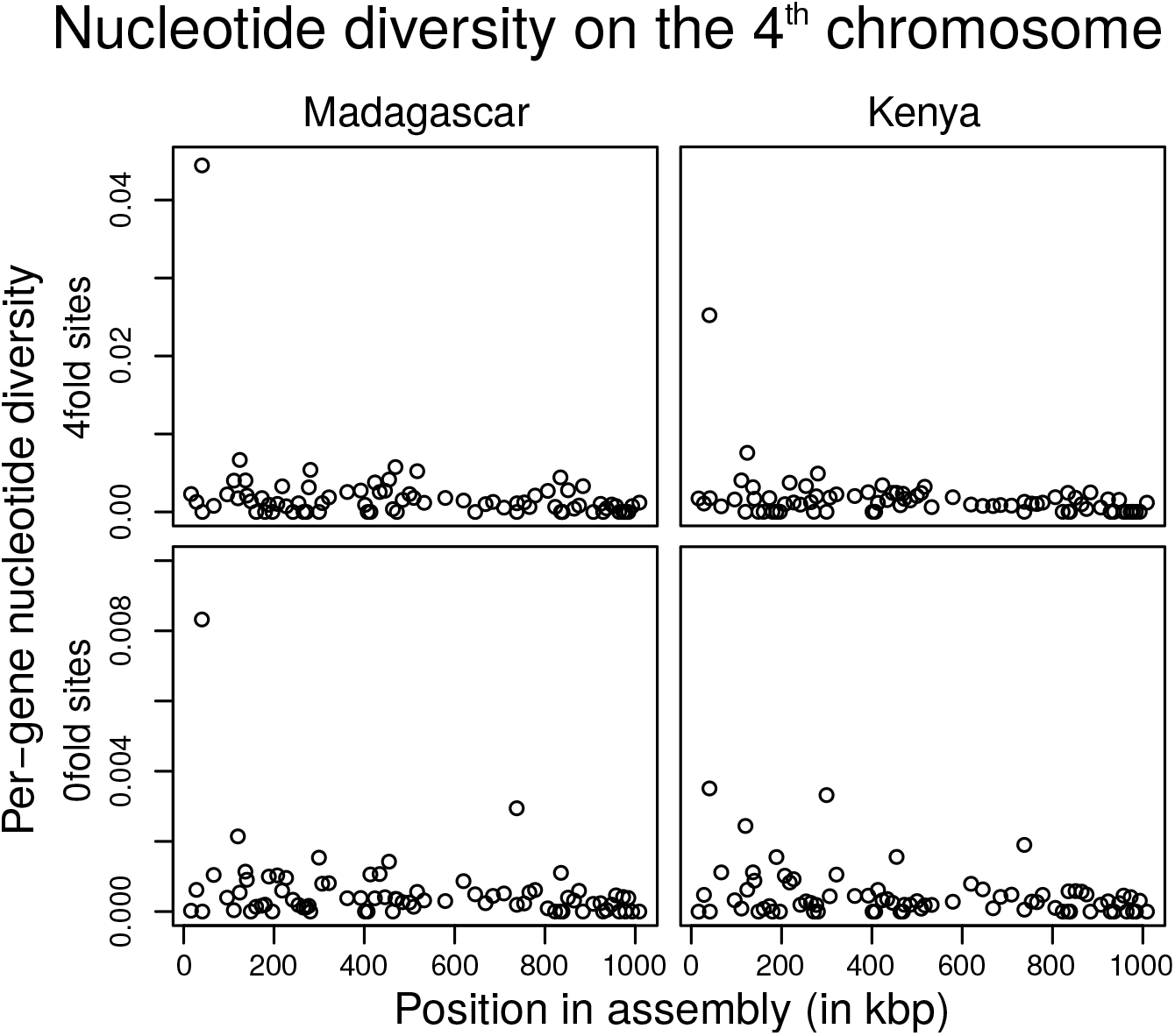
Related to section: Linkage Disequilibrium Patterns and a Further Test for AOD

Supplemental Table S1. Related to Figure 1, Figure 2, and to the first section of the Results and Discussion. Metaanalysis of delta theta in non-crossover regions of other *Drosophila* species.

Supplemental Table S2. Haplotypes around the high-diversity region on the *Drosophila simulans* 4th chromosome. Related to section “Linkage Disequilibrium Patterns and a Further Test for AOD”

Supplemental Table S3. Binned population genetic statistics from two populations of *Drosophila simulans*. Related to figures 1 and S1

Supplemental Table S4. Results of forwards-in-time simulations with parameters relevant to *Drosophila* populations. Related to figure 2.

## STAR Methods

### LEAD CONTACT AND MATERIALS AVAILABILITY

Hannes Becher

### EXPERIMENTAL MODEL AND SUBJECT DETAILS

The results described in here are based on whole genome sequences of isofemale lines of *D. simulans*, established from fertilized females collected in Kenya and Madagascar by William Ballard. These were maintained in the laboratory of Peter Andolfatto for several generations, and subsequently inbred by brother-sister mating in the Charlesworth laboratory. The details of the breeding procedures and sequencing methods are described in [7].

### METHOD DETAILS

#### Sampling and sequencing

We downloaded raw read data in fastq format for 21 isofemale Madagascan genomes and 18 Kenyan genomes from the European Nucleotide Archive (study accession numbers: PRJEB7673; PRJNA215932), as in [7]. We mapped the reads to version 2.02 of the *D. simulans* genome (FlyBase release 2017_04) [48] using BWA MEM [49], then sorted, merged and marked duplicates on the resulting BAM files using Picard Tools version 2.8.3 (https://broadinstitute.github.io/picard/). We called variants separately for each individual line using the HaplotypeCaller tool from GATK version 3.7 [50] with the options – emitRefConfidence, BP_RESOLUTION and -max-alternate-alleles 2, then made per-chromosome VCF files for the whole population using the GATK v3.7 tools combineGVCFs and genotypeGVCFs. All the scripts necessary for downloading the fastq files and calling variants are available at https://github.com/benja-mincjackson/dsim_variant_pipeline_ref_v2.02.git. We defined sites as 4-fold degenerate in all transcripts using information from the gff format annotation of the *D. simulans* genome v2.02 (available from ftp://ftp.flybase.net/genomes/Drosophila_simulans/dsim_r2.02_FB2017_04/gff/dsim-all-r2.02.gff.gz).

#### Data analyses

In the absence of a high-density genetic map for *D. simulans*, we used the mapping data of [31] for *D. melanogaster*. As described in [10], the *D. melanogaster* crossing over (CO) rates were smoothed by Loess regressions against the physical positions of markers in the genome. Each *D. simulans* gene was assigned the CO rate per megabase in female meiosis of the corresponding position in *D. melanogaster*, using an alignment of the genome sequences of *D. simulans* (v2.02) and *D. melanogaster* (v5.57) [48]. Following [10], the absence of recombinational exchange between homologous chromosomes in *Drosophila* males was taken into account by multiplying the autosomal CO rates by one-half, and the X linked rates by two-thirds. All CO rates presented below have been corrected in this way.

For genomic regions with non-zero CO rates, genes were grouped into ten equally-sized bins. For the autosomes (A), there were approximately 730 genes per bin, and approximately 180 genes per bin for the X chromosome (X). All *D. simulans* genes that were aligned to *D. melanogaster* scaffold heterochromatin, or to regions defined as lacking CO in [10], were excluded before binning. These non-CO genes were then binned by chromosome (2: 86, 3: 54, X (tip): 23, X (centromere): 33), and were used for the analyses of the non-CO regions described below.

Chromosome arm 3R, which contains a large inversion relative to *D. melanogaster* [51], was excluded from our analyses. The genetic maps of the X chromosome appear to be similar in the two species [52] so that we used the *D. melanogaster* CO rate estimates as a proxy for those in *D. simulans*. For the analyses of chromosome 2 and 3L, the binning procedure assumes the rank order of the CO rate of a gene to be related to its physical position in the same way as in *D. melanogaster*, even though the absolute CO rates per megabase differ between the species: *D. simulans* has higher CO rates, especially in the low CO rate regions near the centromere and telomere [52]. Non-parametric rank correlations were therefore used to assess statistical relationships between the summary statistics for the autosomal polymorphism data and CO rates.

Levels of polymorphisms per gene were measured by estimates of mean pairwise nucleotide site diversity (π) [8] and Watterson’s theta (θ_w_) [9] for fourfold and zero-fold sites, distinguished here by subscripts 4 and 0. The extent of skew of the folded site frequency spectra (SFS) at fourfold and zero-fold sites for a CO bin was assessed in various ways: the proportion of singleton variants (*P*_*S*_), the mean value of Tajima’s *D* statistic (*D*_T_) across genes in a bin [53], and the relative difference between the mean values of θ_w4_ and π (Δθ_w4_ = 1 − π /θ_w4_) for the genes in a bin. This is related to the Δπ statistic of [54], but has the opposite sign and is not multiplied by Watterson’s correction factor for sample size [9]. An excess of rare variants over the equilibrium neutral expectation for mutation and drift under the infinite sites model [35] is indicated by an excess of *P*_*S*_ over the theoretical value of the probability of observing derived variants present at frequencies of 1/*n* and 1 − 1/*n* in a sample of *n* alleles [9], by a negative value of *D*_T_, and by a positive value of Δθ_w_. The last statistic has the advantage of being independent of the sample size at mutation-drift equilibrium with constant *N*_*e*_ and is used in the figures presented in the text. Results for the other statistics are shown in the Supplemental Figure S1.

LD was computed between pairs of 4-fold sites as the square of Pearson’s correlation coefficient, *r*, between the allelic states of the sites [55]. Pairs of sites were then grouped by distance into bins of 10 kbp and the mean *r*^2^ was computed for each bin, discarding bins over 5 × 10^5^ bp apart, since these contained very few datapoints.

### Simulations

Forwards-in-time simulations were carried out with SLiM v2.6 [56]. The values of the deterministic parameters of mutation, selection and recombination were chosen on the basis of estimates from *D. melanogaster*, as described by [12, 20]. For the simulations, these values were multiplied by (1.33 × 10^6^)/*N*, where *N* is the number of diploid individuals used in the simulations, to ensure that the products of *N* and these parameters correspond approximately to the values for a real *Drosophila* population [12]; this rescaling should conserve the properties of the evolutionary process, with some exceptions discussed in [12]. Population sizes of 2500, 10000 and 25000 diploid individuals were simulated, with a 1:1 sex ratio. The genome consisted of 69 intergenic regions of 4 kb each, and 70 genes plus 4kb of non-coding sequence at both ends of the genome. Each gene was made up of a 5’ and a 3’ UTR of 190bp and 280bp, respectively, and five coding exons of 300 bp separated by four 100bp introns, resulting in a total gene length of 2370 bp and a total chromosome length of 449900bp. This provides an approximate model of the *Drosophila* fourth chromosome, the best-studied non-CO genomic region [57].

A constant unscaled per-basepair mutation rate of 4.5 × 10^−9^ was assumed. To speed up the simulations, no mutations were allowed in the intergenic regions, but CO and gene conversion were allowed at uniform rates over the whole region. Following [12], all intronic mutations were neutral. For simulations that included both background selection (BGS) and selective sweeps (SSWs), UTR mutations were either deleterious or beneficial, with a probability of being beneficial of *p*_*u*_ = 9.04 × 10^−4^ (see Table 1). Coding regions were assumed to consist of 30% 4-fold degenerate sites and 70% 0-fold sites. 30% of new mutations were thus neutral; of the remainder, most were deleterious, with a probability of *p*_*a*_ = 2.21 × 10^−4^ that a mutation at such a site was advantageous [12]. In addition, simulations were run with SSWs only, or with BGS only. Some purely neutral simulations were also run for the purpose of testing. For these configurations, the appropriate mutation types were replaced by neutral ones.

Gamma distributions of selection coefficients were used for the deleterious mutations, and exponential distributions for the beneficial mutations (see Methods Table 1). Since the main question investigated here is the possibility of associative overdominance effects in genomic regions with low CO rates, which requires partial recessivity of deleterious mutations [5], we adjusted the scaled selection coefficients shown in Methods Table 1 by a factor of 1/(2*h*) for runs with dominance coefficient *h* that differed from 0.5, in order to maintain the same heterozygous fitness effects of mutations. In large, randomly mating populations of the type used here, the fates of new mutations are largely determined by their heterozygous fitness effects [16, 58]. The value of *h* should thus not greatly affect the rates of fixation of deleterious and beneficial mutations, or their probability distributions at segregating sites, provided that it is sufficiently far from zero.

Following [12], the standard CO rate in female meiosis was set to 2 × 10^−8^, and a range of values below this, including no CO, was simulated. The rate of initiation of gene conversion (GC) tracts in female meiosis was set to 4 × 10^−8^, double the observed rate for *D. melanogaster* [32], to compensate for the fact that gene conversion in SLiM v2.6 assumes that tracts are initiated in only one direction. The mean tract length was 440 bp, and individual values were drawn from a geometric distribution. No recombination was allowed in males, so that the evolutionarily effective autosomal recombination rate is half the assigned rate.

The simulations were run for 35,000 (= 14*N*) generations and a sample of 20 genomes was taken every *N* generations. The first 20,000 (= 8*N*) generations were treated as a burn-in period, during which the population approached mutation-selection-drift equilibrium, except for runs with zero crossing over, where equilibrium was approached very quickly because of strong effects of selection at linked sites. A few simulations were carried out with larger values of *N* (10,000 and 25,000) in the absence of crossing over. For these, we used the same burn-in and run times, because the simulations reached equilibrium much sooner than with crossing over. Summary statistics were calculated from the state of the population after the simulation finished, and fixations of new mutations were recorded if they arose after the burn-in period.

The large population sizes were used to check whether the rather strong selection against deleterious mutations with a population size of 2,500 caused a very strong haplotype structure in the absence of CO. This can occur because of the build-up of pseudo-overdominance between complementary haplotypes of the type 1010… /0101…, where 1 denotes the wild-type state at a site, and 0 denotes the mutant state [59–62]. However, there were relatively small differences in outcomes for the different population sizes, and even the smallest population size showed no evidence for such an effect (see Figure 3).

Selection parameters used in the simulations with *h* = 0.5.

Selection parameters used in the simulations with $h = 0.5$
| Genomic region | Fitness effect | Proportion of different types of mutations in a region | Distribution of homozygous selection coefficients (scaled by $2N$) |
| --- | --- | --- | --- |
| Coding | Beneficial | $0.7p_a$<br>($p_a = 2.21 \times 10^{-4}$) | Exponential (mean = 250) |
|  | Neutral | 0.3 |  |
| | Deleterious | $0.7(1 - p_a)$ | Gamma (mean = 2000, shape = 0.3) |
| UTR | Beneficial | $p_u = 9.04 \times 10^{-4}$ | Exponential (mean = 213) |
| | Deleterious | $1 - p_u$ | Gamma (mean= 110, shape = 0.3) |
| Intron | Neutral | 1 |  |
| Intergenic | Neutral | 1 |  |

For simulations with other $h$ values, the mean selection coefficients are multiplied by $0.5/h$ .

### QUANTIFICATION AND STATISTICAL ANALYSIS

For the analysis of the *D. simulans* population genetic statistics, genes were binned by crossover rate (10 bins of approx. 730 genes for autosomes, 10 bins of approx. 180 genes for the X chromosome) as detailed in Supplementary Table S1. Per-bin means, weighted by the number of relevant sites per gene, were analysed with Spearman rank correlations for the autosomes, and with least-squares regression models (containing a quadratic term) for the X. Both methods were implemented in R. The *p* values reported are the overall values for the rank correlations, and for the quadratic terms in the X models.

The confidence intervals for the empirical and simulated data were generated via bootstrapping using the R package “boot”. For the empirical data, bootstrapping was carried out on the level of genes per bin (see Table S1). All simulations were run with 20 replicates, which provide the basis for bootstrapping. We report the 95% confidence intervals of the mean based on 1000 bootstrap replicates. For analyses of linkage disequilibrium, we used the same approach for the medians of the *r*^2^ statistic described above.

The decay of linkage disequilibrium with pairwise distance between sites was assessed using Mantel tests as implemented in the R package “vegan”. The value of the Mantel statistic and the associated p-values are reported in the legend of figure 3.

## DATA AND CODE AVAILABILITY

The original data are available from the European Nucleotide Archive, accession numbers PRJEB7673 and PRJNA215932. The scripts developed here are available on GitHub.

## KEY RESOURCES TABLE

Submitted separately as requested.

## Notes

#### Summary of Updates

Author list corrected

